# Neural substrates of cognitive impairment in a NMDAR hypofunction mouse model of schizophrenia and rescue by risperidone

**DOI:** 10.1101/2023.01.16.524241

**Authors:** Cristina Delgado-Sallent, Thomas Gener, Pau Nebot, Cristina López-Cabezón, M. Victoria Puig

## Abstract

NMDAR hypofunction is a pathophysiological mechanism relevant for schizophrenia. Acute administration of the NMDAR antagonist phencyclidine (PCP) induces psychosis in patients and animals while subchronic PCP (sPCP) produces cognitive dysfunction for weeks. We investigated the neural correlates of memory and perceptual impairments in mice treated with sPCP and the rescuing abilities of the atypical antipsychotic drug risperidone administered daily for two weeks. We recorded neural activities in the medial prefrontal cortex (mPFC) and the dorsal hippocampus (dHPC) during memory acquisition, short-term, and long-term memory in the novel object recognition test and during auditory perception and mismatch negativity (MMN) and examined the effects of sPCP and sPCP followed by risperidone. We found that the information about the new object and its short-term storage were associated with mPFC→dHPC high gamma connectivity whereas long-term memory retrieval depended on dHPC→mPFC theta connectivity. sPCP impaired short-term and long-term memory, which was associated with increased mPFC and decreased dHPC neural network activities, and disrupted mPFC-dHPC connectivity. Risperidone rescued the memory deficits and attenuated hippocampal desynchronization. sPCP also impaired auditory perception and its neural correlates (evoked potentials and MMN) in the mPFC, which were also ameliorated by risperidone. Our study suggests that during NMDAR hypofunction the mPFC and the dHPC disconnect possibly underlying cognitive impairment in schizophrenia, and that risperidone targets this circuit to ameliorate cognitive abilities in patients.

## INTRODUCTION

N-methyl D-aspartate receptor (NMDAR) hypofunction is a pathophysiological mechanism found in schizophrenia patients that can be replicated in rodent models by using pharmacological agents that block NMDAR, for example phencyclidine or ketamine (Lee and Zhou, 2019). Sub-chronic administration of phencyclidine (sPCP, also known as angel dust) to rodents mimics cognitive symptoms in schizophrenia for several months. That is, it impairs executive functions such as recognition memory, cognitive flexibility, and sensorimotor gating, including mismatch negativity (MMN)(Rajagopal et al., 2014; Hamilton et al., 2018b; Lee and Zhou, 2019). Moreover, exposure to PCP reduces the density of PV-expressing GABAergic interneurons in the prefrontal cortex (PFC) and the hippocampus (HPC) (Abdul-Monim et al., 2007) of the animals. This is similar to what is observed in post mortem tissue from patients with schizophrenia (Lewis et al., 2005; Konradi et al., 2011; Kaar et al., 2019), further validating sPCP as a suitable model of schizophrenia. The reduction in PV-expressing neuron populations impair perisomatic inhibition of pyramidal neurons that likely contributes to a diminished gamma synchronization that is required for most cognitive functions (Lewis et al., 2005). However, a comprehensive neurophysiological characterization of prefrontal-hippocampal circuits following sPCP treatment is missing.

Atypical antipsychotic drugs are effective in reducing the positive symptoms of schizophrenia patients while showing modest amelioration of the negative and cognitive symptoms. The neural substrates underlying these complex behavioural effects are poorly understood, which makes the development of better treatments a difficult endeavour. In rodents, atypical antipsychotic drugs attenuate some of the behavioural effects induced by NMDAR antagonists, including sPCP-induced cognitive dysfunction (Grayson et al., 2007). They enhance dopamine efflux in the cortex and the HPC, affecting less the limbic system, whereas classical neuroleptics show the opposite effect. These actions are mediated by the affinity of atypical antipsychotic drugs for serotonin receptors, especially the 1A (5-HT_1A_R) and the 2A (5-HT_2A_R) subtypes, which have a widespread expression in the brain and modulate the dopaminergic, serotonergic, glutamatergic, and GABAergic systems. Risperidone is one of the most prescribed atypical antipsychotic drugs. It targets mainly D2R and 5-HT_2A_R and is effective at treating the positive symptoms while also ameliorating certain aspects of cognitive symptoms in humans and rodents, including executive function, attention, learning, and memory (Grayson et al., 2007; Houthoofd et al., 2008; Rajagopal et al., 2014; Baldez et al., 2021).

We have recently reported that acute PCP exerts strong influences on prefrontal-hippocampal neural dynamics in freely moving mice. The psychosis-like states produced by PCP were associated with hypersynchronisation of the medial PFC (mPFC), desynchronization of the dorsal HPC (dHPC), and disrupted mPFC-dHPC circuit connectivity. Acute risperidone reduced cortical hypersynchronisation but had limited efficacy in restoring hippocampal synchronization and circuit connectivity (Delgado-Sallent et al., 2022). In healthy mice, acute risperidone had an inhibitory effect on the circuit, reducing spiking activity and theta-gamma oscillations in both areas, and it also disrupted the circuit’s connectivity (Gener et al., 2019). Here, we investigated the neural correlates of memory and perceptual deficits in the sPCP model of schizophrenia and their rescue by chronic risperidone, experimental conditions closer to the clinical settings.

## METHODS AND MATERIALS

### Animals

Experiments were performed in C57BL/6J male mice (*n* = 51) that were 2 to 3 months old at the start of the experiments. Mice were housed under conditions of controlled temperature (23 ±1°C) and illumination (12 hours light/dark cycle). All procedures were conducted in compliance with EU directive 2010/63/EU and Spanish guidelines (Laws 32/2007, 6/2013 and Real Decreto 53/2013) and were authorized by the Barcelona Biomedical Research Park (PRBB) Animal Research Ethics Committee and the local government.

### Surgeries

Mice were induced with a mixture of ketamine/xylazine and placed in a stereotaxic apparatus. Anesthesia was maintained with continuous 0.5-4% isoflurane. Small craniotomies were drilled above the medial PFC and HPC. Several micro-screws were screwed into the skull to stabilize the implant, and the one on top of the cerebellum was used as a general ground. Three tungsten electrodes, one stereotrode and one single electrode, were implanted in the prelimbic region of the medial PFC (mPFC) and two more were implanted in the CA1 area of the dorsal HPC (dHPC). The electrodes were positioned stereotaxically in the prelimbic cortex (AP: 1.5, 2.1 mm; ML: ± 0.6, 0.25 mm; DV: −1.7 mm from bregma) and the CA1 region (AP: −1.8 mm; ML: −1.3 mm; DV: −1.15 mm). Neural activity was recorded while the electrodes were being lowered down inside the brain to help locate the CA1 region. In addition, three reference electrodes were implanted in the corpus callosum and lateral ventricles (AP: 1, 0.2, −1; ML: 1, 0.8, 1.7; DV: −1.25, −1.4, −1.5, respectively). The electrodes were made with two twisted strands of tungsten wire 25 μm wide (Advent, UK) and were held together using heat insulation. At the time of implantation, the electrodes had an impedance from 100 to 400 kΩ and were implanted unilaterally with dental cement. Electrode wires were pinned to an adaptor to facilitate their connection to the recording system. After surgery, animals were allowed at least one week to recover during which they were extensively monitored and received both analgesia and anti-inflammatory treatments. Additionally, animals were handled and familiarized with the implant connected to the recording cable. After the experiments ended, the electrode placements were confirmed histologically by staining the brain slices with Cresyl violet (Figure 1B). Electrodes with tips outside the targeted areas were discarded from data analyses.

**FIGURE 1.**
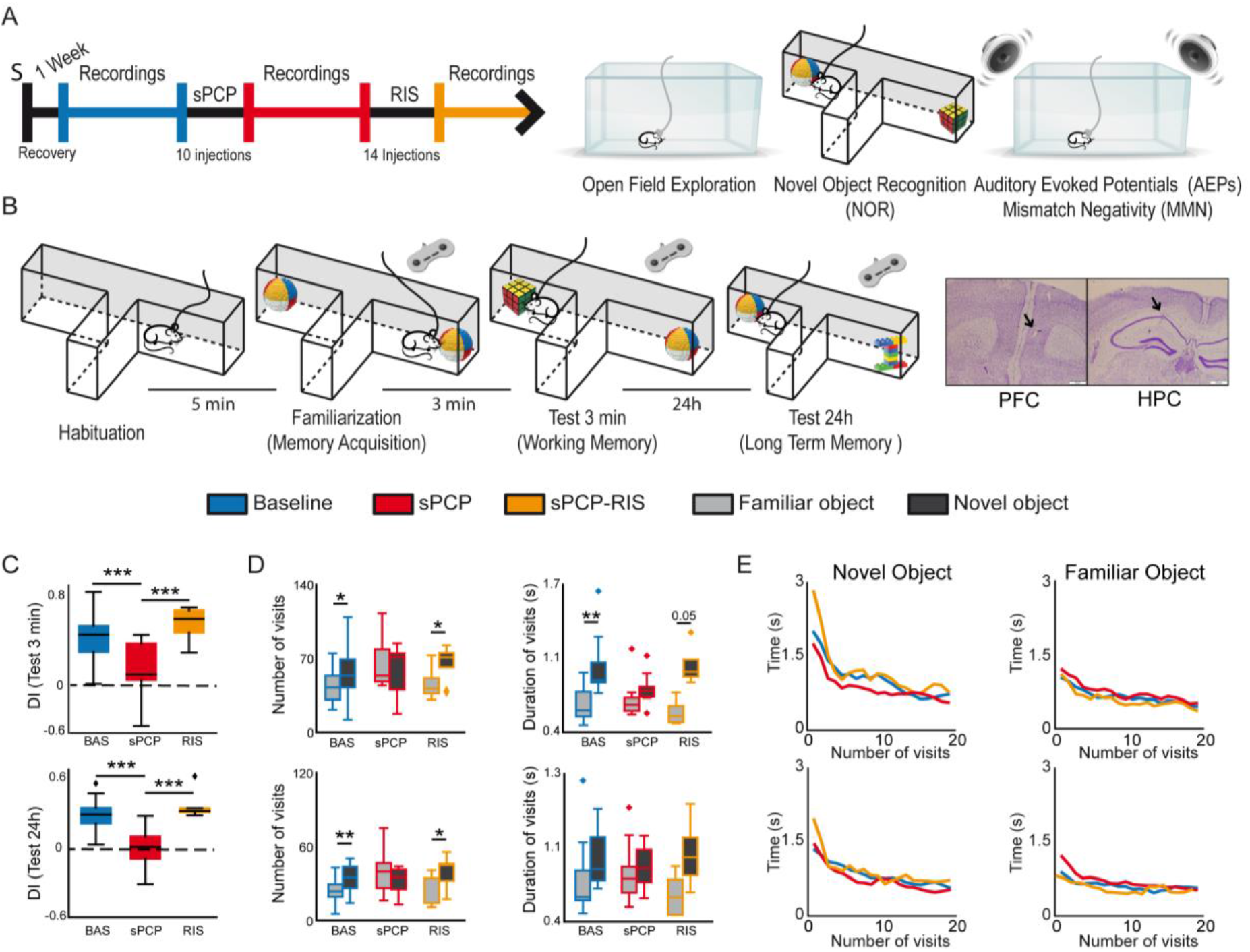
Experimental design and main behavioural results. **(A)** Experimental protocol and behavioural tests used. S indicates the date of surgery to implant electrodes. **(B)** The novel object recognition test (NOR) in detail. Right and left buttons on a joystick were pressed for the duration of each visit to timestamp the information into the recording files. Representative examples of histological verification of electrode locations within the prelimbic PFC and the CA1 area of the HPC. We recorded neural activities during the surgical implantations to increase the chances of implanting the electrodes within the target areas. At the end of the experiment, before euthanasia, a gentle current was passed through the electrodes to mark the recording areas (arrows). **(C)** Discrimination indices (DIs) during the 3min (STM) and the 24h (LTM) tests. DIs decreased significantly after sPCP and recovered with chronic risperidone (STM, LTM: *F*_(2,6)_ = 16.64, 15.15, *p* = 0.004, < 0.0005, one-way ANOVA). **(D)** Number and mean duration of visits to familiar and novel objects during the STM and LTM tests. After sPCP, mice visited both objects evenly by increasing the number and duration of visits to familiar objects, reflecting poor recognition memory (*F*_(1,13)_ = 15.12, 5.2, *p* = 0.002, 0.037, one-way ANOVA). **(E)** The duration of the visits decreased within a session as the mice became less interested in the objects ([baseline STM, LTM]: *F*_(2,26)_ = 3.52, 3.56, *p* = 0.005, 0.0006; one-way ANOVA with novel and familiar objects combined). A sharp decrease in duration occurred during the first 5 visits to both objects in the two tests, and this was more pronounced for novel objects (first 5 visits to novel objects vs. first 5 visits to familiar objects; [STM, LTM]: *F*_(2,26)_ = 36.03, 11.57,*p* < 0.0005, 0.002; one-way ANOVA). These behaviours were not overtly disrupted by sPCP or risperidone.

### Behavioral tests

#### Resting states

Recordings during quiet wakefulness were performed in an open field box (50×40×20 cm) for 30 minutes. We harnessed the accelerometer signals integrated within the Intan RHD2132 amplifiers to precisely monitor general mobility of mice. We quantified the variance of the instantaneous acceleration module (ACC; variance(Root square[X^2^, Y^2^, Z^2^])) that was maximum during exploration and decreased as the animals were in quiet alertness (Gener et al., 2019; Delgado-Sallent et al., 2022). Low mobility was determined by a defined threshold in the output of the accelerometer signals and normal movement was defined as above that threshold. The animals typically rested for brief periods between 2 to 10 consecutive seconds.

#### Novel object recognition test (NOR)

We quantified recognition memory in a custom-designed T-maze as previously reported (Alemany-González et al., 2020, 2022). The maze was made of aluminum with wider and higher arms than the standard mazes (8 cm wide x 30 cm long x 20 cm high). The maze was shielded and grounded for electrophysiological recordings and was placed on an aluminum platform. The novel-familiar object pairs were previously validated as in (Gulinello et al., 2018). The arm of the maze where the novel object was placed was randomly chosen across experiments. The task consisted of a habituation phase, familiarization phase, short-term memory test (STM), and long-term memory test (LTM), each lasting ten minutes (Figure 1B). During the habituation phase, the animals explored the maze without objects. Five minutes later, mice were put back in the maze where two identical objects had been placed at the end of the lateral arms for the familiarization phase. Three minutes and twenty-four hours later, a familiar object and a new object were placed in the maze for the STM and LTM tests, respectively. Memory acquisition was investigated by comparing the initial and the last visits to the two identical items in the familiarization phase; memory retrieval and novelty seeking were investigated during the initial visits to familiar and novel objects, respectively, during the STM and LTM tests. Exploratory events were timestamped into the recording files by sending TTL pulses to the acquisition system. This allowed us to precisely quantify the number and duration of the visits. We used this information to estimate the neurophysiological correlates of memory performance. The duration of the initial visits and the visits to the novel objects were typically longer than the last visits of the sessions and to the familiar objects (Figure 1E). Because some of the mathematical tools employed are sensitive to the time window used, we concatenated different visits until the accumulated time reached 5 seconds. We note that familiarization was considered valid when mice visited the objects for at least ten seconds, as in previous studies (Alemany-González et al., 2020, 2022). The exact number of visits used for each analysis was the following: [Familiarization phase] All conditions combined, first: 3.91±0.17, last: 4.86±0.05; BAS, first: 3.81±0.25, last: 4.81±0.1; sPCP, first: 4.25±0.19, last: 5; RIS, first: 3.54±0.45, last: 4.73±0.14; [Three minute test] Novel objects, all conditions combined, 3.24±0.2; BAS: 3.07±0.25, sPCP: 3.92±0.25, RIS: 2.29±0.52; Familiar objects, all conditions combined, 4.29±0.13; BAS: 4.5±0.16, sPCP: 4.07±0.25, RIS: 4.29±0.36; [24h test] Novel objects, all conditions combined, 3.83±0.12; BAS: 3.84±0.21, sPCP: 3.89±0.12, RIS: 3.7±0.26; Familiar objects, all conditions combined, 4.47±0.1; BAS: 4.47±0.15, sPCP: 4.57±0.19, RIS: 4.3±0.15.

#### Auditory evoked potential (AEP) and mismatch negativity (MMN) tests

The auditory tests were conducted consecutively in the home cage located inside a soundproof box. We used the Python library *simpleaudio* to generate the sounds in custom scripts that synchronized the sound with the electrophysiological recordings via an EIB board. Mice were habituated to the environment for 5 minutes. Next, 100 consecutive clicks separated by 10 seconds were presented to the mice for a total of 8 minutes. A click consisted of 15 ms of white noise (‘5ms_whitenoise.wav’ function in the *simpleaudio* Python library). We next used an auditory oddball paradigm to measure MMN. Mice were presented with a series of standard tones (6 or 8 kHz) in which a target tone (8 or 6 kHz) was presented randomly in 75 to 25 % proportions, respectively. The frequencies of the standard and target tones were switched after 500 trials in a flip-flop design (Hamilton et al., 2018a). Tones lasted 10 ms and were presented with 500 ms intertrial intervals. The protocol consisted in the presentation of 1000 tones, lasting around 10 minutes.

### Pharmacology

The doses used were: phencyclidine hydrochloride (PCP; Sigma-Aldrich) 10 mg/kg, 5+5 days; risperidone (RIS; Sigma-Aldrich) 0.5 mg/kg, 14 consecutive days. PCP was administered subcutaneously (SC) and risperidone intraperitoneally (IP). The following pharmacological groups were investigated: sPCP (n = 21, sPCP-RIS (*n* = 9), and their corresponding saline controls: SAL (*n* = 7), sPCP-SAL (*n* = 7), SAL-SAL (*n* = 7; Supplementary Figure 1).

### Neurophysiological recordings and data analyses

All the recordings were implemented with the multi-channel Open Ephys system at 0.1-6000 Hz and a sampling rate of 30 kHz. Recorded signals from each electrode were filtered offline to extract local field potentials (LFPs) and multi-unit activity (MUA).

#### Oscillatory activity

To obtain LFPs, recorded signals were detrended, notch-filtered and decimated to 1kHz offline. The frequency bands considered for the band-specific analyses included: delta (2-5 Hz), slow theta (4-8 Hz), theta (8-12 Hz), low gamma (30-48 Hz), high gamma (52-100 Hz), and HFOs (100-200 Hz). Power spectral density results were calculated using the multi-taper method. Spectrograms were constructed using consecutive Fourier transforms. Phase-amplitude coupling (PAC) was measured following the method described in (Onslow et al., 2011). The parameters used were: phase frequencies = [0, 15] with 1 Hz step and 4 Hz bandwidth, amplitude frequencies = [10, 250] with 5 Hz step and 10 Hz bandwidth. Phase-amplitude coupling quantification results were obtained by averaging the values of selected areas of interest in the comodulograms. The exact frequencies used for each analysis are described in the figure legends. Directionality of signals between areas (PFC→HPC and HPC→PFC) was calculated with the phase slope index (PSI) with a Python translation of MATLAB’s data2psi.m (epleng = 60s, segleng = 1s) as in (Nolte et al., 2008). PSI measures were shuffle-corrected. The surrogate analyses were performed by randomizing the data of two pairs of channels, one in PFC and one in HPC, using all the combinations of different pairs of channels. The data were shuffled across time series and across pair of channels 1000 times to obtain the correspondent PSI shuffle. Next, to remove chance effects, the randomized data were averaged and subtracted from the original data. To establish significance, a *t*-test was performed between the original data and its shuffle (Puig and Miller, 2015; Lancaster et al., 2018).

#### Spiking activity

MUA was estimated by first subtracting the raw signal from each electrode with the signal from a nearby referencing electrode to remove artifacts resulting from the animal’s movement. Then, continuous signals were filtered between 450-6000 Hz with Python and thresholded at −3 sigma standard deviations with Offline Sorter v4 (Plexon Inc). We estimated MUA in one-minute non-overlapping windows. Spike-LFP coupling was estimated with the pairwise phase consistency (PPC) method, which is an unbiased parameter to determine the degree of tuning of the neurons’ firing to ongoing network activity at specific frequencies (Vinck et al., 2011, 20; Zorrilla de San Martín et al., 2020). PPC was determined using the phases of spikes from MUA in 25 second epochs, only considering epochs with at least 250 spikes. We used the instantaneous module of the raw x, y and z signals from the accelerometer (Acc) to evaluate the effects of the drugs on general mobility of mice as in previous studies by our group (Gener et al., 2019; Alemany-González et al., 2020, 2022).

### Statistical Analysis

Data are represented in boxplots (*seaborn* function in Python) where the median and the quartiles are shown. We used paired *t*-tests, one-WAY ANOVAs to compare behavioural and neurophysiological measures within animals (baseline-sPCP-RIS), and mixed ANOVAS to compare behavioral and neurophysiological measures between pharmacological groups (baseline-sPCP-SAL). Two-WAY ANOVAs were used to compare the effects of the sPCP-RIS group with their respective saline controls (sPCP-SAL, SAL-SAL). Paired *t*-tests were used to assess differences between early and late visits to objects (familiarization test), and between visits to novel and familiar objects (3-minute and 24h tests). We used Sidak’s correction *post hoc* tests in the ANOVAs. We employed Pearson correlations to identify associations between neurophysiological measures and DIs. Statistical analyses were implemented in Python with the Pingouin statistical package (Vallat, 2018).

## RESULTS

### Chronic risperidone prevents sPCP-induced working and long-term memory deficits

We administered subchronically to mice the NMDAR antagonist phencyclidine (10 mg/kg SC 5+5 days; sPCP group) to model cognitive impairment in schizophrenia. We investigated the neural substrates of the well-established memory and perceptual deficits of this model and their subsequent rescue by chronic risperidone (0.5 mg/kg IP for 14 consecutive days; sPCP-RIS group). We recorded neural activities in the prelimbic medial PFC and CA1 region of the dorsal HPC, brain regions relevant for cognitive processing and the pathophysiology of schizophrenia (Sigurdsson and Duvarci, 2016). The animals’ behaviours and neural activities were characterized during baseline, after sPCP, after risperidone (sPCP-RIS group; Figure 1A), and their corresponding saline controls (SAL, sPCP-SAL, and SAL-SAL groups; Supplementary Figure 1).

We first examined short-term memory (STM) and long-term memory (LTM) abilities of the different pharmacological groups assessed via the NOR task (Figure 1B), a well validated memory test that relies on the mice’ innate instinct to explore novel objects and depends on hippocampal-prefrontal circuits (Warburton and Brown, 2015; Alemany-González et al., 2020; Wang et al., 2021). Consistent with previous studies, discrimination indices (DIs) for novel versus familiar objects ([time visiting the novel object - time visiting the familiar object] / total exploration time) were positive for all the mice during baseline in the 3-minute and 24h memory tests (DIs = 0.35 ± 0.05 and 0.29 ± 0.04; *n* = 7, 8 mice, respectively). That is, mice exhibited good STM and LTM at the beginning of the experiment. After sPCP, mice explored novel and familiar objects evenly (DIs = 0.01 ± 0.06 and −0.06 ± 0.06; baseline vs. sPCP: *p* < 0.0005, paired *t*-test) indicating poor recognition memory. This was not observed in controls with saline (SAL group, *n* = 4 mice). Chronic risperidone rescued both STM deficits (DI = 0.42 ± 0.1; *n* = 4 mice) and LTM deficits (DI = 0.37 ± 0.07; *n* = 6 mice; Figure 1C), but not their corresponding saline controls (sPCP-SAL group, *n* = 7 mice; Supplementary Figure 1).

We next analyzed the number and duration of the visits to each object per session. During baseline and risperidone conditions, mice visited the novel items on more occasions than the familiar items ([STM: baseline, risperidone]: *p* = 0.019, 0.038; [LTM: baseline, risperidone]: *p* = 0.003, 0.047; paired *t*-tests) and the visits lasted longer, particularly during the STM test ([baseline, risperidone]: *p* < 0.005, 0.056). In contrast, after sPCP the mice visited both objects evenly by increasing the number and duration of visits to familiar objects, behaving as if they were novel (Figure 1D). This indicated poor recognition memory. The duration of the visits decreased within a session as the mice became less interested in the objects, both novel and familiar. A sharp decrease in duration occurred during the first 5 visits to the objects, mainly to the novel objects, in the two tests and subsequently the duration of the visits declined slowly. These prolonged initial visits did not occur when the mice visited the familiar objects. These behaviors were not overtly disrupted by sPCP or risperidone (Figure 1E).

### Risperidone partially rescues sPCP-induced disruptions of theta and gamma synchronization in prefrontal-hippocampal circuits during quiet alertness

We first aimed to gain insight into the neural dynamics of healthy prefrontal-hippocampal circuits (mPFC-dHPC) during quiet alertness and their impact by sPCP and risperidone. To mitigate the effects of the hyperlocomotion produced by sPCP (Mouri et al., 2012; Castañé et al., 2015) we analysed neural activities during resting states, brief 3-second episodes of low behavioural activity while the mice explored an open field (see Methods). As we reported previously (Delgado-Sallent et al., 2022), during resting states healthy mice exhibited a strong circuit synchronization at theta frequencies (4-12 Hz): theta oscillations were prominent in the mPFC and there were robust theta oscillations, theta-gamma and spike-theta coupling in the dHPC (Figure 2A-C).

**FIGURE 2.**
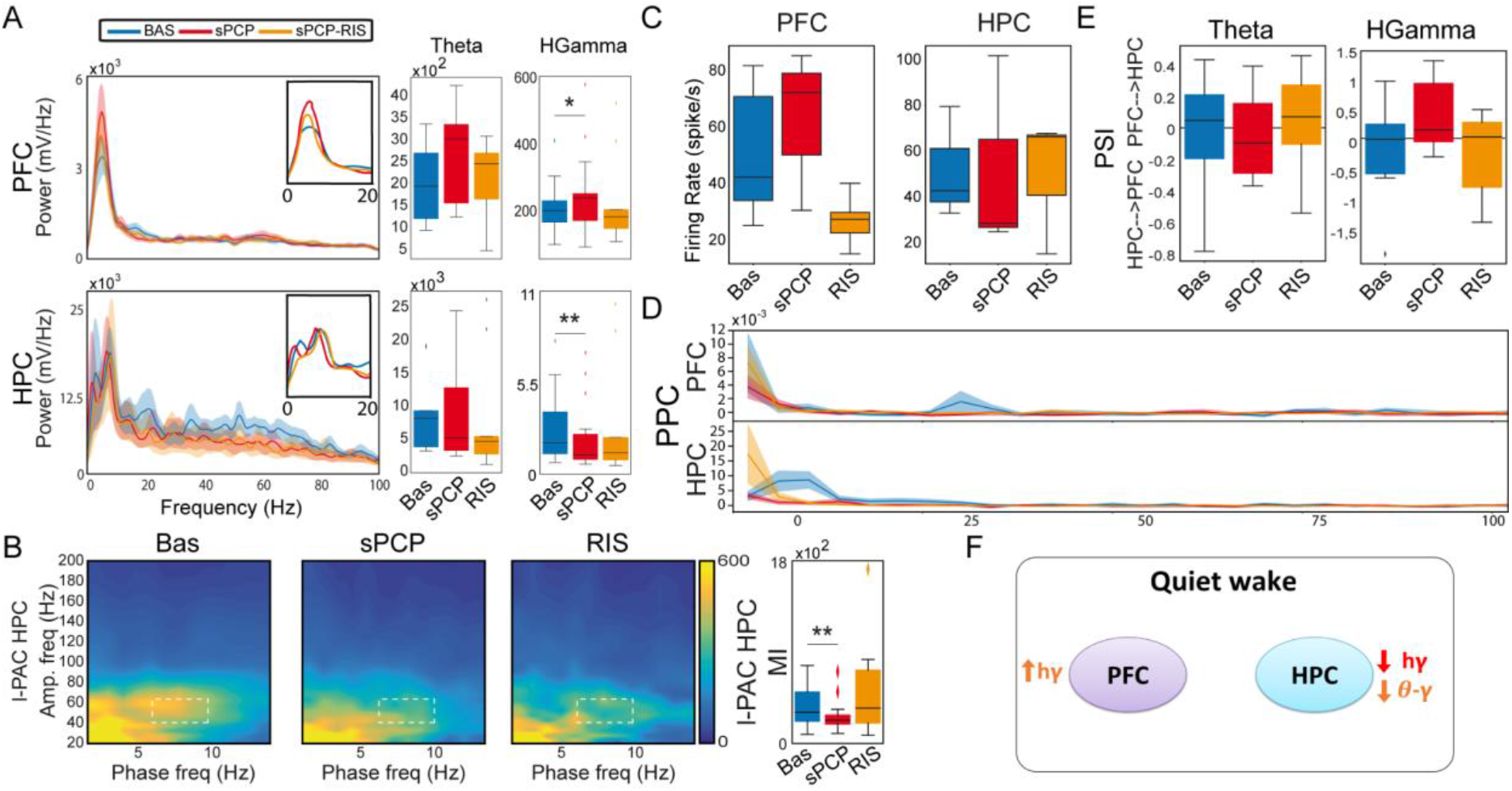
sPCP disrupted theta and gamma synchronization in mPFC-dHPC circuits during quiet alertness. Some of these alterations were ameliorated by risperidone. **(A)** sPCP increased high gamma power in the mPFC (baseline vs. sPCP: *p* = 0.047, paired *t*-test; *n* = 21 mice) and decreased high gamma power in the dHPC (*p* = 0.004). Risperidone rescued aberrant prefrontal, but not hippocampal, power. **(B)** sPCP weakened local and inter-regional theta-gamma coupling (6-10 Hz with 40-60 Hz; l-PAC; *p* = 0.008; ir-PAC PFC_phase_-HPC_amp_; *p* = 0.001) that were partially restored by risperidone. (**C**) sPCP promoted the spiking activity of neuron populations (multi-unit activity or MUA) in the mPFC that was reduced by risperidone (*F*_(2,6)_ = 4.58, *p* = 0.047; one-way ANOVA). **(D)** sPCP disrupted the coupling of spikes to theta oscillations in the dHPC that was restored by risperidone (*F*_(2,6)_ = 19.4, *p* = 0.002; one-way ANOVA). Risperidone also boosted spike-delta coupling in the HPC (3-6 Hz, *F*_(2,6)_ = 5.39, *p* = 0.046). Spike-LFP coupling was estimated via the pairwise phase consistency method (PPC). **(E)** sPCP did not significantly affect the directionality of theta and high gamma signals within the circuit, but some tendencies were observed in the high gamma band. Circuit directionality was estimated via the phase slope index (PSI).

sPCP disrupted theta and gamma rhythms within mPFC-dHPC circuits that ameliorated with risperidone (*n* = 21 mice). First, sPCP increased high gamma (52-100 Hz) power in the mPFC that was reduced by risperidone. Concomitantly, sPCP decreased high gamma power in the dHPC, however this was not rescued by risperidone (Figure 2A). In addition, sPCP weakened intrinsic theta-gamma coupling both locally in the dHPC and inter-regionally that were partially restored by risperidone (Figure 2B). We detected strong correlations between the concomitant reductions of hippocampal high gamma power and the two forms of theta-gamma coupling both during baseline ([HPC, circuit] Pearson’s R = 0.73, 0.69, *p* < 0.0005) and after sPCP (R = 0.81, 0.76, *p* < 0.0005). These correlations put forward a key role of hippocampal high gamma oscillations in the generation of theta-gamma coupling within the circuit. Moreover, sPCP tended to increase the firing rate of neuron populations (MUA) in the mPFC that was reduced by risperidone (*n* = 5 mice; Figure 2C). In the dHPC, sPCP did not change the firing rate of neurons but disrupted the coupling of spikes to ongoing theta oscillations (*n* = 4 mice). Risperidone was unable to restore the spike-theta synchronization and it actually increased the locking of spikes to delta oscillations (< 5 Hz; Figure 2D). No major changes in power, cross-frequency coupling, and spike-LFP coupling were observed at other frequencies in either region. Finally, the directionality of signals within the circuit was not overtly affected by sPCP or risperidone (Figure 2E). These sPCP-induced alterations were not present in saline controls (SAL group, *n* = 7 mice; Supplementary Figure 2). Together, these findings suggested that risperidone partially improved the disrupted theta and gamma synchronization caused by sPCP in mPFC-dHPC circuits (Figure 2F).

### Risperidone partially restores the neural correlates of sPCP-induced poor recognition memory in prefrontal-hippocampal circuits

We first aimed to understand the neural mechanisms underlying memory acquisition in healthy animals (*n* = 9 mice). We compared the neurophysiological signals during the familiarization phase (Figure 1B) between the initial visits to the two (identical) objects, which imply novelty seeking, and the last visits of the session, when the animals had just acquired a new memory about the objects. More specifically, we compared neurophysiological biomarkers recorded during the first five seconds versus the last five seconds of the total accumulated time of the visits, regardless of object (i.e., located in the right or left arms) or number of visits. This was a good trade-off because familiarization was considered valid when mice visited the objects for at least ten seconds, as in previous studies (Alemany-González et al., 2020, 2022). On average, the 5-second epochs included 3.91 ± 0.2 early visits and 4.86 ± 0.06 late visits (see Methods). During normal memory acquisition, theta power (8-12 Hz) increased in the mPFC and decreased in the dHPC (early vs. late visits; *p* = 0.012, 0.024, paired *t*-test; Figure 3A), whereas local and inter-regional theta-gamma coupling seemed unchanged (Figure 3B). We note that changes in theta power were not simply due to differential locomotion of mice during a session, as the animals’ mobility was low while they were visiting the objects both at the beginning and at the end of the sessions. We further investigated whether any flow of information emerged within the circuit upon memory acquisition. We focused on theta and gamma frequencies based on previous work from our group (Alemany-González et al., 2020, 2022). We found that mPFC→dHPC high gamma signals tended to emerge during the late visits (Figure 3C). Therefore, memory acquisition was associated with increases and decreases of theta power in the mPFC and the dHPC, respectively, and mPFC→dHPC high gamma signals.

**FIGURE 3.**
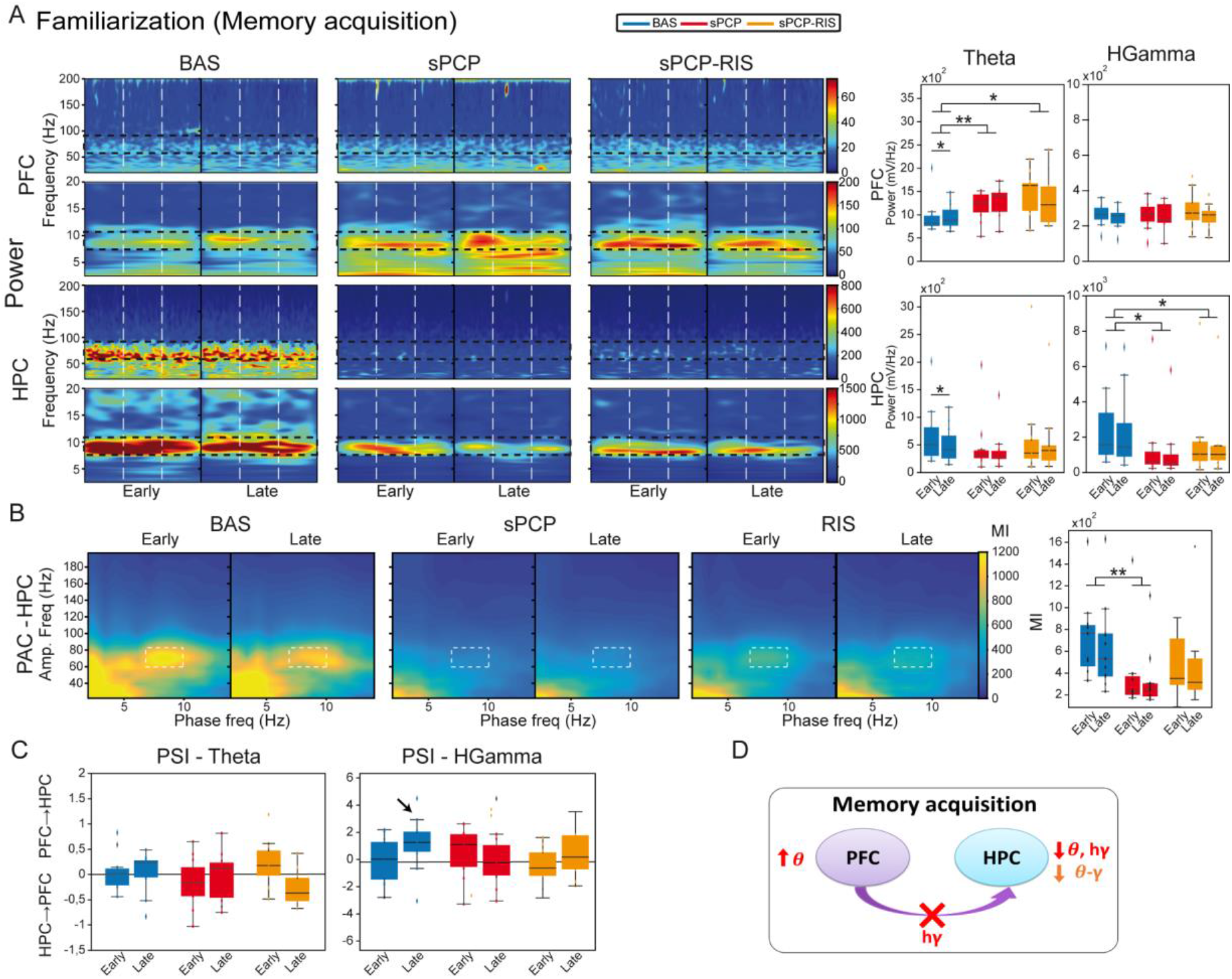
Neural substrates of memory acquisition in the NOR task and effects of sPCP and risperidone. **(A)** Prefrontal theta power increased in healthy animals during the late visits to the objects. sPCP increased non-specifically theta power in the mPFC (baseline vs. sPCP: *F*_(1,10)_ = 7.46, *p* = 0.021; one-way ANOVA combining early and late visits) and reduced theta and high gamma power in the HPC (baseline vs. sPCP: *F*_(1,12)_ = 4.81, 9.59, *p* = 0.053, 0.011), disrupting the normal neural dynamics of memory acquisition. Risperidone did not rescue these power changes. For visualization purposes only, we show the second preceding the initiation of the visit, the visit itself (which typically lasted less than one second) and the subsequent second (the first line thus indicates the start of the button-press on the joystick). **(B)** Intrinsic dHPC theta-gamma coupling (7-10 Hz with 60-80 Hz) did not change with memory acquisition. sPCP disrupted theta-gamma coordination that was partially rescued by risperidone ([l-PAC] *F*(1,12) = 37.68,*p* < 0.0005; one-way ANOVA). Similar results were obtained for inter-regional theta-gamma coupling ([ir-PAC] *F*_(1,12)_ = 41.09, *p* < 0.0005; data not shown). **(C)** In healthy mice, mPFC→dHPC high gamma signals tended to occur during the late visits to the objects (PSI vs. shuffle, *p* = 0.069; marked with an arrow). sPCP disrupted these flows of information (baseline vs. sPCP, *p* = 0.096; PSI vs. shuffle, *p* = 0.49) that were not restored by risperidone (PSI vs. shuffle, *p* = 0.38). The PSI was shuffle-corrected. **(D)** Proposed neural mechanism for memory acquisition and effects of sPCP and risperidone. In red, changes produced by sPCP, in orange sPCP-induced deviations ameliorated by risperidone.

sPCP augmented non-specifically theta power (8-12 Hz) in the mPFC and decreased it in the dHPC. In addition, high gamma power also declined in the dHPC after sPCP (Figure 3A). None of the power changes were consistently recovered by risperidone, although tendencies were detected in the dHPC. Local and inter-regional theta-gamma did not change during memory acquisition. However, as above, and consistent with the reduction of gamma power in the dHPC, both types of theta-gamma coupling weakened after sPCP (differences with SAL controls; Supplementary Figure 2). After risperidone, theta-gamma coupling was partially restored (Figure 3B). Finally, sPCP disrupted the mPFC→dHPC high gamma signals detected during late visits to the objects that were partially corrected by risperidone (Figure 3C). Overall, we found that memory acquisition was encoded by progressive changes in theta power in the mPFC and the dHPC and mPFC→dHPC high gamma signals that emerged during the late visits. sPCP disrupted these biomarkers and further weakened theta-gamma coordination that were partially restored by risperidone (Figure 3D).

We next investigated the neural substrates of STM (*n* = 7 mice). We compared the neurophysiological signals recorded during the first five seconds of visits to the novel and familiar objects during the 3-minute tests (Figure 1B). On average, these included 3.24 ± 0.2 and 4.29 ± 0.13 visits to novel and familiar objects, respectively (Figure 1E). Animals with DIs over 0.2 during the baseline were used in these analyses. Theta power (Figure 4A) and theta-gamma coupling (Figure 4B) did not differ during the visits to novel and familiar objects. However, we detected a mPFC→dHPC flow of information at high gamma frequencies during the visits to familiar objects (Figure 4C), similar to memory acquisition. This directionality of signals seemed relevant for memory processing as it correlated strongly with memory performances (DI with PSI: Pearson’s R = 0.7, *p* = 0.026). Similar again to memory acquisition, sPCP increased theta waves in the PFC and reduced high gamma power in the dHPC (Figure 4A). Correspondingly, both local and inter-regional theta-gamma coupling were also reduced (Figure 4B). Risperidone ameliorated the power and coupling alterations produced by sPCP in the HPC, but the measures were not fully recovered. The mPFC→dHPC high gamma signals detected during familiar visits when the animals were healthy were also disrupted by sPCP and were not recovered by risperidone (Figure 4C).

**FIGURE 4.**
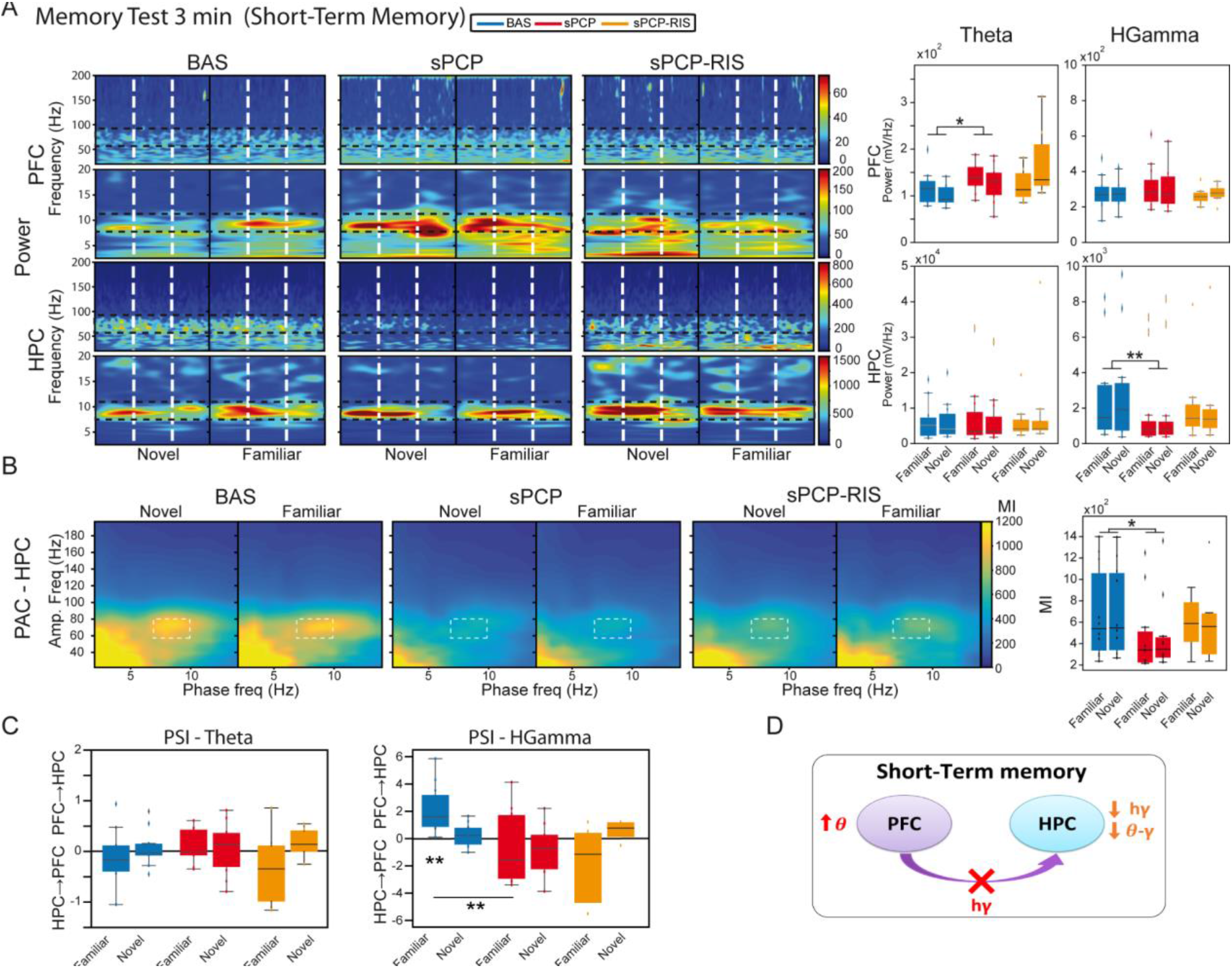
Neural substrates of working memory during the 3-minute tests and effects of sPCP and risperidone. **(A)** Theta and high gamma oscillations were similar during the visits to familiar and novel objects in both regions. sPCP increased theta waves in the mPFC non-specifically interfering with the normal theta dynamics (*F*_(1,7)_ = 8.26 *p* = 0.024; mixed ANOVA with object (familiar vs novel) and treatment (baseline vs sPCP) as factors). sPCP also reduced dHPC high gamma power (*F*_(1(,12)_ = 8.37, *p* = 0.023; differences with sPCP-SAL controls: *F*_(1,20)_ = 6.44, *p* = 0.022) that were partially rescued by risperidone. For visualization purposes only, we show the second used for the analyses, the second prior and the second later separated by dashed lines, the first line thus indicating the start of the button-press on the joystick. **(B)** Intrinsic hippocampal theta-gamma coupling (7-10 Hz with 60-80 Hz) was similar during visits to familiar and novel objects. sPCP disrupted theta-gamma coordination that was partially rescued by risperidone ([l-PAC] *F*_(1,12)_ = 10.36, *p* < 0.009; differences with sPCP-SAL controls). Similar results were obtained for inter-regional theta-gamma coupling ([ir-PAC] *F*_(1,12)_ = 16.38, *p* = 0.002, data not shown; differences with sPCP-SAL controls). **(C)** In healthy mice, mPFC→dHPC high gamma signals were detected during the visits to the familiar objects (PSI vs. shuffle, *p* = 0.004). sPCP disrupted this flow of information (baseline vs. sPCP, *p* = 0.008; PSI vs. shuffle, *p* = 0.19) that was not restored by risperidone (PSI vs. shuffle, *p* = 0.24) The PSI was shuffle-corrected. **(D)** Proposed neural mechanism for STM and effects of sPCP and risperidone. In red, changes produced by sPCP, in orange sPCP-induced deviations ameliorated by risperidone.

We further investigated the neural substrates of LTM (*n* = 8 mice). We compared the neurophysiological signals recorded during the first five seconds of visits to the novel and familiar objects during the 24h memory tests (Figure 1B). On average, these included 3.83±0.12 and 4.47±0.1 visits to novel and familiar objects, respectively (Figure 1E). As above, only animals with DIs over 0.2 during baseline were used. Similar to STM, we did not detect changes in power or theta-gamma coupling between visits to novel and familiar objects in either region (Figure 5A,B). Furthermore, dHPC→mPFC theta signals were detected during the visits to familiar objects and correlated inversely with the DIs (Pearson’s R = −0.72 *p* = 0.01; Figure 5C). As above, sPCP decreased dHPC high gamma power when the animals visited both objects (Figure 5A). Also as above, both local and inter-regional theta-gamma coupling were reduced by sPCP (Figure 5B). Like during STM processes, risperidone partially rescued sPCP-reduced gamma power and theta-gamma coupling in the dHPC. However, it was unable to restore theta and high gamma dHPC→mPFC signals disrupted by sPCP (Figure 5C).

**FIGURE 5.**
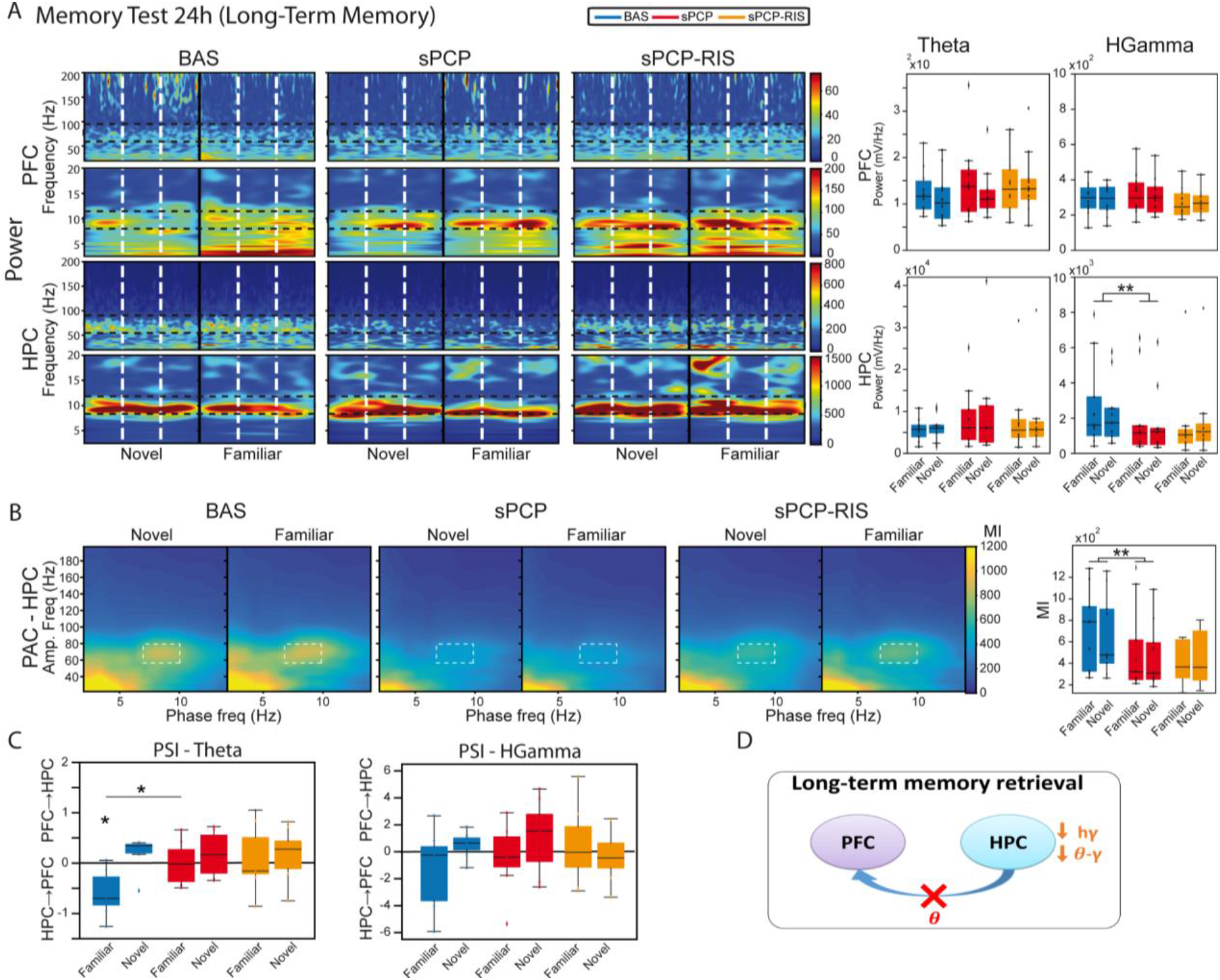
Neural substrates of LTM during the 24h tests and effects of sPCP and risperidone. **(A)** Power in the mPFC and the dHPC was similar during visits to familiar and novel objects. sPCP reduced high gamma power in the dHPC non-specifically (*F*_(1,12)_ = 15.33, *p* = 0.006; differences with SAL controls). As above, risperidone partially rescued hippocampal gamma power. For visualization purposes only, we show the second used for the analyses, the second prior and the second later separated by dashed lines, the first line thus indicating the start of the button-press on the joystick. **(B)** Intrinsic hippocampal theta-gamma coupling (7-10 Hz with 60-80 Hz) was similar during visits to familiar and novel objects. sPCP disrupted theta-gamma coordination that was partially rescued by risperidone ([l-PAC] *F*_(1,12)_ = 15.96,*p* < 0.003; differences with sPCP-SAL controls). Similar results were obtained for inter-regional theta-gamma coupling ([ir-PAC] *F*_(1,12)_ = 33.19,*p* < 0.0005; differences with sPCP-SAL controls; data not shown). **(C)** In healthy mice, dHPC→mPFC theta signals were detected during the visits to the familiar objects (PSI vs. shuffle, *p* = 0.031). sPCP disrupted this flow of information (baseline vs. sPCP, *p* = 0.024; PSI vs. shuffle, *p* = 0.94) that was not rescued by risperidone (PSI vs shuffle, *p* = 0.64). The PSI has been shuffle-corrected. **(D)** Proposed neural mechanism for LTM and effects of sPCP and risperidone.

Together, these results unraveled a contribution of theta and gamma signals within mPFC-dHPC circuits to normal memory acquisition, STM and LTM processes within the context of object recognition memory. sPCP increased theta power in prefrontal microcircuits, disrupted high gamma rhythms in the dHPC and the theta-gamma coupling associated with it, regardless of brain state and cognitive task. That is, sPCP-induced neurophysiological alterations were observed across different days. Risperidone partially rescued some of these disturbances, hippocampal gamma oscillations and corresponding theta-gamma coupling in a more consistent way, suggesting a preferential action on this brain region with respect to the mPFC and the circuit’s connectivity.

### Risperidone improves impaired auditory processing and its neural correlates in sPCP-treated mice

We finally investigated the effects of sPCP and risperidone on the neural substrates of auditory attention and perception. More specifically, we examined alterations in auditory evoked potentials (AEPs) and mismatch negativity (MMN), as both biomarkers are highly translational between patients and rodent models of schizophrenia (Amann et al., 2010; Nagai et al., 2013; Javitt and Sweet, 2015; Modi and Sahin, 2017). Briefly, mice were placed in a cage surrounded by a sound enclosure with four speakers. The animals were presented with standard auditory stimuli first and subsequently the oddball paradigm in 30-minute sessions during baseline, after sPCP and after risperidone treatments as above. Only animals that conducted the three experiments were included in the analyses. Neural activities were recorded both in the mPFC and the dHPC, but hippocampal responses were inconsistent and highly variable between animals, therefore we only present data collected from the mPFC.

The AEP protocol consisted of 100 trials of white noise, each tone lasting 15 ms with an inter-trial interval of 10s (Figure 6A). During baseline, we detected cortical AEPs in 82.3% of the trials (response ratio: [number of AEPs / number of trials]). The response ratio decreased after sPCP to 64.7%, but not after saline (Supplementary Figure 3). Risperidone partially rescued the response ratio to 71.8% (Figure 6B). Concomitantly, sPCP-treated mice showed abnormal AEPs with respect to the baseline, specifically the P2 and P3 peaks, analogous to the P200 and P300 peaks in humans. sPCP attenuated the spiking activity associated with the P2 component (40-70 ms) whereas the P3 component (200-300 ms) was reduced both in amplitude and corresponding spiking activity. Risperidone ameliorated the alterations associated with both components (Figure 6C).

**FIGURE 6.**
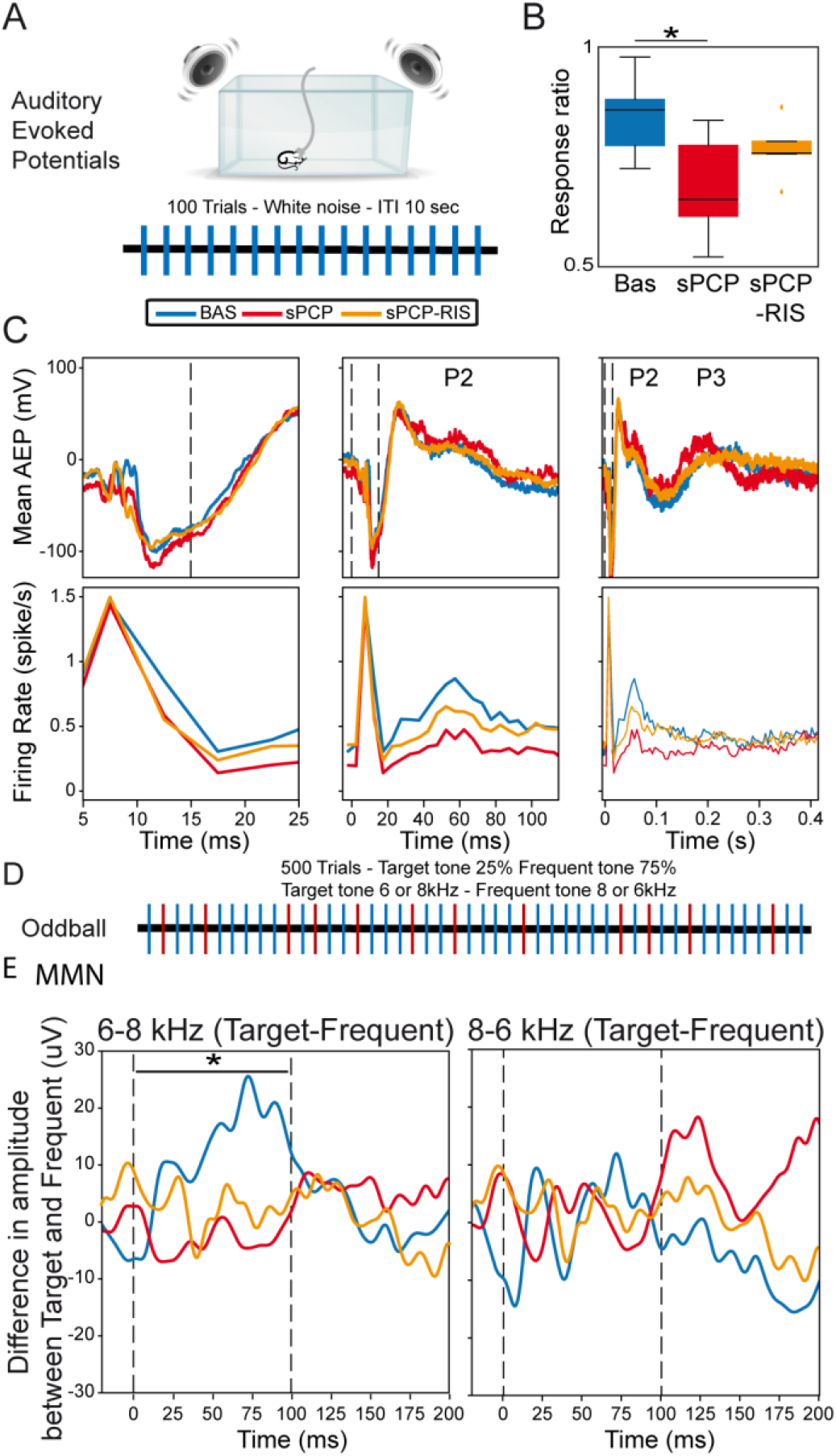
Neural substrates of auditory processing in the prelimbic cortex and effects of sPCP and risperidone. **(A)** AEP protocol. **(B)** Ratio of AEP responses detected in the PFC after the auditory stimuli. The ratio decreased in sPCP-treated animals ([baseline vs. sPCP, sPCP vs. saline]: *p* = 0.024, 0.024, paired and unpaired *t*-test, respectively) and was partially recovered by risperidone. **(C)** Mean AEP and corresponding spiking activity (multi-unit firing rates) at three different timescales. P2 (40-70 ms) was associated with less spiking activity after sPCP ([baseline vs. sPCP, sPCP vs. saline as above]: *p* = 0.013, 0.068), whereas P3 (200-300 ms) was reduced in amplitude and spiking activity ([baseline vs. sPCP, sPCP vs. saline as above]: amplitude: *p* = 0.015, 0.055; firing rate: *p* = 0.024, 0.06). Risperidone increased the spiking activity associated within P2 with respect to sPCP (F_2,10_ = 2.49, *p* = 0.076) and augmented the amplitude and spiking activity associated with P3 component ([amplitude, firing rate]: *p* = 0.065, 0.097). **(D)** Protocol used for the oddball paradigm to assess mismatch negativity (MMN). **(E)** MMN was detected during the presentation of the 6-8 KHz target-frequent combination (left) but not the 8-6 kHz combination (right). Shown is the subtraction of target-frequent responses. MMN was absent in the sPCP-treated group (baseline vs. sPCP area under the curve; *p* = 0.021, paired *t*-test) and not restored by risperidone. Vertical dashed lines mark the start and end of tone presentation.

We finally examined the MMN using a passive oddball paradigm. Mismatch negativity is a component of event-related potentials that reflects preattentive auditory sensory memory. It emerges following deviant auditory stimuli and has been well characterized in patients with schizophrenia. We examined whether sPCP-treated mice exhibited reduced MMN and the rescuing abilities of risperidone. Mice were presented with 500 tones that differed in frequency (6 kHz or 8 kHz) in 25-75% ratios (target and standard stimuli, respectively; Figure 6D). During baseline and in SAL mice, MMN emerged when the 6/8 kHz target-standard tone combination was presented, but not with the 8/6 kHz combination. The MMN was abolished by sPCP and not restored by risperidone (Figure 6E and Supplementary Figure 3). Together, risperidone attenuated the behavioural and neurophysiological alterations associated with the late components of the AEPs, but failed to rescue the MMN.

## DISCUSSION

We found that subchronic PCP impaired STM, LTM, and auditory perception in mice that were rescued by risperidone administered daily for two weeks. The behavioural deficits were accompanied by disrupted dHPC-mPFC neural dynamics that attenuated after risperidone. These findings suggest that risperidone targets this circuit to elicit its beneficial actions on cognitive abilities in patients with schizophrenia.

The memory impairments produced by sPCP and their rescue by risperidone are in consonance with previous studies reporting that sPCP disrupts recognition memory in rodents that is sensitive to antipsychotic medication (Castañé et al., 2015; Rajagopal et al., 2016; Cadinu et al., 2018). Despite the novel object recognition task being extensively used as a screening tool for memory abilities in biomedicine, an understanding of its underlying neural mechanisms was missing. Thus, we first investigated whether and how mPFC-dHPC circuits encoded memory acquisition, STM, and LTM in this task. Our findings unravelled a relevant role for the circuit’s communication in the encoding of the NOR task during baseline, with limited contribution of local power and theta-gamma coupling. mPFC→dHPC high gamma signals emerged during memory acquisition and the visits to familiar objects in the 3-minute memory test, and were strongly associated with STM performance (R = 0.7). Later, dHPC→mPFC theta signals were detected during the visits to familiar objects in the 24h memory test, which also correlated with LTM retrieval (R = 0.72). We note that the mPFC sends direct afferents to the dHPC in the mouse whereas dHPC→mPFC pathways use the nucleus reuniens of the thalamus as a relay (Sigurdsson and Duvarci, 2016).

Our findings suggest that the information about the new object (acquisition) and its short-term storage were encoded by direct mPFC→dHPC connectivity whereas long-term memory retrieval depended on dHPC→mPFC indirect connectivity. This hypothesis is in line with previous studies showing that novel experiences initiate molecular signalling in the PFC that subsequently travel to the HPC (Takehara-Nishiuchi, 2020), where memories are stored (Eichenbaum, 2004). A key contribution of the circuit’s connectivity to memory processing was evidenced further by the fact that sPCP impaired STM and LTM and disrupted dHPC-mPFC communication during the three phases of the NOR task. Previous studies from our group have consistently detected that dHPC→mPFC signals are relevant for long-term memory retrieval in wild-type mice of a different genetic background and altered in a mouse model of intellectual disability (Alemany-González et al., 2020, 2022), further supporting the findings presented here. Conflicting results exist on whether recognition memory for objects requires the PFC (Spanswick and Dyck, 2012; Morici et al., 2015). Recent studies using optogenetic interrogation however implicate the mPFC in this type of memory and, in fact, identify HPC→mPFC theta coupling as a major neural mechanism involved (Wang et al., 2021; Chao et al., 2022).

The differential effects of sPCP on prefrontal and hippocampal microcircuits may explain the disruption of the flow of information within this pathway. sPCP “disconnected” the circuit by causing opposite effects in the two brain regions: it increased neural synchronization in the mPFC (enhanced theta and gamma power) and desynchronized neural networks in the dHPC (reduced theta, gamma power, and theta-gamma coupling). Prior studies have demonstrated that acute NMDAR hypofunction desynchronizes neural activity in the HPC of rodents, including oscillatory and cross-frequency coupling (Caixeta et al., 2013). Remarkably, we recently reported comparable effects following an acute administration of PCP that generated psychosis-like states in mice (Delgado-Sallent et al., 2022). In that study, acute risperidone attenuated aberrant cortical hypersynchronisation but was unable to restore hippocampal desynchronization. This likely reflects risperidone’s efficacy in containing psychosis but its inefficiency in ameliorating cognitive abilities in the short term. While acute risperidone has proven to be effective in restoring cognitive deficits in the sPCP mouse model of schizophrenia (Grayson et al., 2007; Meltzer et al., 2011; Cadinu et al., 2018), it was important to assess the effects of a chronic treatment that mimicked more realistically the prescription of antipsychotic drugs to patients. Here, the injection of risperidone for two weeks rescued STM and LTM impairments and attenuated the neural activity reductions observed in the dHPC, albeit it was unable to restore cortical hypersynchronisation or the connectivity of the circuit. However, it is plausible that dampening cortical hyperactivity by more prolonged administration of risperidone restores the circuit’s communication. Therefore, chronic medication with risperidone may indeed favour healthier dHPC-mPFC neural dynamics in patients with schizophrenia, accounting for their better performance in executive function, attention, learning, and memory (Houthoofd et al., 2008; Baldez et al., 2021).

Furthermore, the findings of this study may help reconcile studies in schizophrenia patients, in which both increases and decreases of gamma oscillations were found to be key biomarkers of the disorder (Uhlhaas and Singer, 2010). Pathological gamma oscillations in the cortex may originate from disinhibition of cortical pyramidal neurons resulting from deficient PV-expressing interneurons (Sigurdsson, 2016; Guyon et al., 2021; Alemany-González et al., 2022). In contrast, sPCP reduces PV-interneuron density in the HPC (Abdul-Monim et al., 2007; Sabbagh et al., 2013) that, in turn, down-regulates pyramidal neuron activity (Korotkova et al., 2010). This may explain the dampening of gamma oscillations and theta-gamma coupling observed in the dHPC of sPCP-treated mice. Consistent with our findings, abnormal gamma synchrony produced by NMDAR hypofunction in both areas has been linked to cognitive impairment, including tasks assessing STM and object recognition (Tort et al., 2009; Korotkova et al., 2010; Carlén et al., 2012). Further experiments however are needed to elucidate the exact neural mechanisms involved in the generation of abnormal gamma oscillations in schizophrenia. Risperidone attenuated sPCP-induced gamma and theta-gamma decreases in the dHPC very consistently. This APD binds to 5-HT_2A_R with high affinity for which it is an inverse agonist. Therefore, it is likely that risperidone modulates gamma synchrony via 5-HT_2A_R-expressing PV interneurons, which are present both in the mPFC and the dHPC (Puig et al., 2010; Puig and Gulledge, 2011; Puig and Gener, 2015).

We finally investigated abnormal auditory processing in sPCP-treated mice, the underlying neural correlates in the frontal cortex, and the rescuing abilities of risperidone. As above, we found that sPCP impaired auditory perception in the mPFC that was attenuated by risperidone. That is, there was a reduction in the number of AEPs after sPCP, which in turn exhibited smaller P2 and P3 late peaks, the mouse components analogous to the P200 and P300 peaks in humans (Amann et al., 2010; Modi and Sahin, 2017). In addition, abnormal P2 and P3 components were associated with decreased spiking activity. Risperidone partially restored the late components of the AEPs, as observed in patients (Umbricht et al., 1999). We note that we did not detect alterations in the early component of AEPs (the P1 and N1), which have been associated with positive symptoms (Galderisi et al., 2014), consistent with the lack of psychotic-like symptoms in sPCP-treated animals. However, electrophysiological recordings in the auditory cortex would be necessary to confirm these results as the auditory cortex has shown more consistent alterations in the early components in patients and rodent models (Wang et al., 2020). sPCP-treated mice also showed reduced MMN, however it was not rescued by risperidone. Reduced amplitude of the P200 and P300 peaks and the MMN are robust findings in patients with schizophrenia (Galderisi et al., 2014; Javitt and Sweet, 2015; Turetsky et al., 2015; Hamilton et al., 2019; Fitzgerald and Todd, 2020; Koshiyama et al., 2020) and have also been reported in NMDAR hypofunction animal models (Ehrlichman et al., 2008, 2009; Amann et al., 2010). Therefore, our findings on sPCP- and risperidone-produced alterations of AEPs and MMN are highly translational and relevant to human studies.

In conclusion, sPCP impaired recognition memory that was associated with increased mPFC, decreased dHPC neural network activities, and disrupted mPFC-dHPC communication. Risperidone rescued the memory deficits and attenuated hippocampal desynchronization. sPCP also impaired auditory attention and its neural correlates in the mPFC, which were also ameliorated by risperidone. Our study strongly suggests that NMDAR hypofunction disconnects the mPFC and the dHPC underlying cognitive impairment in schizophrenia, and that risperidone targets this circuit to ameliorate cognitive abilities in patients.

## ACKNOWLEDGEMENTS

We thank Jose A. Garrido and Amanda B. Fath for scientific insight into this study and Patricia Ruiz for administrative support. This study was financed by grants SAF2016-80726-R and PID2019-104683RB-I00 to MVP funded by MCIN/AEI/10.13039/501100011033 and by ERDF “A way of making Europe”. CD-S was supported by a FI AGAUR predoctoral fellowship from the Catalan government (Generalitat de Catalunya grant number 2018 FI_B_00112).

## REFERENCES

Abdul-Monim, Z., Neill, J. C., and Reynolds, G. P. (2007). Sub-chronic psychotomimetic phencyclidine induces deficits in reversal learning and alterations in parvalbumin-immunoreactive expression in the rat. J. Psychopharmacol. Oxf. Engl. 21, 198–205. doi: 10.1177/0269881107067097.

Alemany-González, M., Gener, T., Nebot, P., Vilademunt, M., Dierssen, M., and Puig, M. V. (2020). Prefrontal-hippocampal functional connectivity encodes recognition memory and is impaired in intellectual disability. Proc. Natl. Acad. Sci. 117, 11788–11798. doi: 10.1073/pnas.1921314117.

Alemany-González, M., Vilademunt, M., Gener, T., and Puig, M. V. (2022). Postnatal environmental enrichment enhances memory through distinct neural mechanisms in healthy and trisomic female mice. Neurobiol. Dis. 173:105841. doi: 10.1016/j.nbd.2022.105841.

Amann, L. C., Gandal, M. J., Halene, T. B., Ehrlichman, R. S., White, S. L., McCarren, H. S., et al. (2010). Mouse behavioral endophenotypes for schizophrenia. Brain Res. Bull. 83, 147–161. doi: 10.1016/j.brainresbull.2010.04.008.

Baldez, D. P., Biazus, T. B., Rabelo-da-Ponte, F. D., Nogaro, G. P., Martins, D. S., Kunz, M., et al. (2021). The effect of antipsychotics on the cognitive performanc of individuals with psychotic *disorders: Network meta-analyses of randomized controlled trials*. Neurosci. Biobehav. Rev. 126, 265–275. doi: 10.1016/j.neubiorev.2021.03.028.

Cadinu, D., Grayson, B., Podda, G., Harte, M. K., Doostdar, N., and Neill, J. C. (2018). NMDA receptor antagonist rodent models for cognition in schizophrenia and identification of novel drug treatments, an update. Neuropharmacology 142, 41–62. doi: 10.1016/j.neuropharm.2017.11.045.

Caixeta, F. V., Cornélio, A. M., Scheffer-Teixeira, R., Ribeiro, S., and Tort, A. B. L. (2013). Ketamine alters oscillatory coupling in the hippocampus. Sci. Rep. 3, 2348. doi: 10.1038/srep02348.

Carlén, M., Meletis, K., Siegle, J. H., Cardin, J. A., Futai, K., Vierling-Claassen, D., et al. (2012). A critical role for NMDA receptors in parvalbumin interneurons for gamma rhythm induction and behavior. Mol. Psychiatry 17, 537–548. doi: 10.1038/mp.2011.31.

Castañé, A., Santana, N., and Artigas, F. (2015). PCP-based mice models of schizophrenia: differential behavioral, neurochemical and cellular effects of acute and subchronic treatments. Psychopharmacology (Berl.) 232, 4085–4097. doi: 10.1007/s00213-015-3946-6.

Chao, O. Y., Nikolaus, S., Yang, Y.-M., and Huston, J. P. (2022). Neuronal circuitry for recognition memory of object and place in rodent models. Neurosci. Biobehav. Rev. 141, 104855. doi: 10.1016/j.neubiorev.2022.104855.

Delgado-Sallent, C., Nebot, P., Gener, T., Fath, A. B., Timplalexi, M., and Puig, M. V. (2022). Atypical, but not typical, antipsychotic drugs reduce hypersynchronized prefrontal-hippocampal circuits during psychosis-like states in mice: Contribution of 5-HT2A and 5-HT1A receptors. Cereb. Cortex 32, 3472–3487. doi: 10.1093/cercor/bhab427.

Ehrlichman, R. S., Gandal, M. J., Maxwell, C. R., Lazarewicz, M. T., Finkel, L. H., Contreras, D., et al. (2009). N-methyl-d-aspartic acid receptor antagonist-induced frequency oscillations in mice recreate pattern of electrophysiological deficits in schizophrenia. Neuroscience 158, 705–712. doi: 10.1016/j.neuroscience.2008.10.031.

Ehrlichman, R. S., Maxwell, C. R., Majumdar, S., and Siegel, S. J. (2008). Deviance-elicited changes in event-related potentials are attenuated by ketamine in mice. J. Cogn. Neurosci. 20, 1403–1414. doi: 10.1162/jocn.2008.20097.

Eichenbaum, H. (2004). Hippocampus: cognitive processes and neural representations that underlie declarative memory. Neuron 44, 109–120. doi: 10.1016/j.neuron.2004.08.028.

Fitzgerald, K., and Todd, J. (2020). Making sense of mismatch negativity. Front. Psychiatry 11, 468. doi: 10.3389/fpsyt.2020.00468.

Galderisi, S., Vignapiano, A., Mucci, A., and Boutros, N. N. (2014). Physiological correlates of positive symptoms in schizophrenia. Curr. Top. Behav. Neurosci. 21, 103–128. doi: 10.1007/7854_2014_322.

Gener, T., Tauste-Campo, A., Alemany-González, M., Nebot, P., Delgado-Sallent, C., Chanovas, J., et al. (2019). Serotonin 5-HT1A, 5-HT2A and dopamine D2 receptors strongly influence prefronto-hippocampal neural networks in alert mice: Contribution to the actions of risperidone. Neuropharmacology 158, 107743–107743. doi: 10.1016/j.neuropharm.2019.107743.

Grayson, B., Idris, N., and Neill, J. (2007). Atypical antipsychotics attenuate a sub-chronic PCP-*induced cognitive deficit in the novel object recognition task in the rat*. Behav. Brain Res. 184, 31–38. doi: 10.1016/j.bbr.2007.06.012.

Gulinello, M., Mitchell, H. A., Chang, Q., Timothy O’brien, W., Zhou, Z., Abel, T., et al. (2018). Rigor and reproducibility in rodent behavioral research. Neurobiol Learn Mem S1074-7427, 30001–7. doi: 10.1016/j.nlm.2018.01.001.

Guyon, N., Zacharias, L. R., Oliveira, E. F. de, Kim, H., Leite, J. P., Lopes-Aguiar, C., et al. (2021). Network asynchrony underlying increased broadband gamma power. J. Neurosci. doi: 10.1523/JNEUROSCI.2250-20.2021.

Hamilton, H. K., D’Souza, D. C., Ford, J. M., Roach, B. J., Kort, N. S., Ahn, K.-H., et al. (2018a). Interactive effects of an N-methyl-d-aspartate receptor antagonist and a nicotinic acetylcholine receptor agonist on mismatch negativity: Implications for schizophrenia. Schizophr. Res. 191, 87–94. doi: 10.1016/j.schres.2017.06.040.

Hamilton, H. K., Perez, V. B., Ford, J. M., Roach, B. J., Jaeger, J., and Mathalon, D. H. (2018b). Mismatch negativity but not P300 is associated with functional disability in schizophrenia. Schizophr. Bull. 44, 492–504. doi: 10.1093/schbul/sbx104.

Hamilton, H. K., Woods, S. W., Roach, B. J., Llerena, K., McGlashan, T. H., Srihari, V. H., et al. (2019). Auditory and visual oddball stimulus processing deficits in schizophrenia and the psychosis risk syndrome: Forecasting psychosis risk with P300. Schizophr. Bull. 45, 1068–1080. doi: 10.1093/schbul/sby167.

Houthoofd, S. A. M. K., Morrens, M., and Sabbe, B. G. C. (2008). Cognitive and psychomotor effects of risperidone in schizophrenia and schizoaffective disorder. Clin. Ther. 30, 1565–1589. doi: 10.1016/j.clinthera.2008.09.014.

Javitt, D. C., and Sweet, R. A. (2015). Auditory dysfunction in schizophrenia: integrating clinical and basic features. Nat. Rev. Neurosci. 16, 535–550. doi: 10.1038/nrn4002.

Kaar, S. J., Angelescu, I., Marques, T. R., and Howes, O. D. (2019). Pre-frontal parvalbumin interneurons in schizophrenia: a meta-analysis of post-mortem studies. J. Neural Transm. Vienna Austria 1996 126, 1637–1651. doi: 10.1007/s00702-019-02080-2.

Konradi, C., Yang, C. K., Zimmerman, E. I., Lohmann, K. M., Gresch, P., Pantazopoulos, H., et al. (2011). Hippocampal interneurons are abnormal in schizophrenia. Schizophr. Res. 131, 165–173. doi: 10.1016/j.schres.2011.06.007.

Korotkova, T., Fuchs, E. C., Ponomarenko, A., von Engelhardt, J., and Monyer, H. (2010). NMDA receptor ablation on parvalbumin-positive interneurons impairs hippocampal synchrony, spatial representations, and working memory. Neuron 68, 557–569. doi: 10.1016/j.neuron.2010.09.017.

Koshiyama, D., Kirihara, K., Tada, M., Nagai, T., Fujioka, M., Usui, K., et al. (2020). Reduced auditory mismatch negativity reflects impaired deviance detection in schizophrenia. Schizophr. Bull. 46, 937–946. doi: 10.1093/schbul/sbaa006.

Lancaster, G., Iatsenko, D., Pidde, A., Ticcinelli, V., and Stefanovska, A. (2018). Surrogate data for hypothesis testing of physical systems. Phys. Rep. 748, 1–60. doi: 10.1016/j.physrep.2018.06.001.

Lee, G., and Zhou, Y. (2019). NMDAR hypofunction animal models of schizophrenia. Front. Mol. Neurosci. 12, 185. doi: 10.3389/fnmol.2019.00185.

Lewis, D. A., Hashimoto, T., and Volk, D. W. (2005). Cortical inhibitory neurons and schizophrenia. Nat. Rev. Neurosci. 6, 312–324. doi: 10.1038/nrn1648.

Meltzer, H. Y., Horiguchi, M., and Massey, B. W. (2011). The role of serotonin in the NMDA *receptor antagonist models of psychosis and cognitive impairment*. Psychopharmacology (Berl.) 213, 289–305. doi: 10.1007/s00213-010-2137-8.

Modi, M. E., and Sahin, M. (2017). Translational use of event-related potentials to assess circuit integrity in ASD. Nat. Rev. Neurol. 13, 160–170. doi: 10.1038/nrneurol.2017.15.

Morici, J. F., Bekinschtein, P., and Weisstaub, N. V. (2015). Medial prefrontal cortex role in recognition memory in rodents. Behav. Brain Res. 292, 241–251. doi: 10.1016/J.BBR.2015.06.030.

Mouri, A., Koseki, T., Narusawa, S., Niwa, M., Mamiya, T., Kano, S., et al. (2012). Mouse strain differences in phencyclidine-induced behavioural changes. Int. J. Neuropsychopharmacol. 15, 767–779. doi: 10.1017/S146114571100085X.

Nagai, T., Tada, M., Kirihara, K., Araki, T., Jinde, S., and Kasai, K. (2013). Mismatch negativity as a *“translatable” brain marker toward early intervention for psychosis: a review*. Front. Psychiatry 4, 115. doi: 10.3389/fpsyt.2013.00115.

Nolte, G., Ziehe, A., Nikulin, V. V., Schlögl, A., Krämer, N., Brismar, T., et al. (2008). Robustly estimating the flow direction of information in complex physical systems. Phys Rev Lett 100, 234101–234101. doi: 10.1103/PhysRevLett.100.234101.

Onslow, A. C. E., Bogacz, R., and Jones, M. W. (2011). Quantifying phase-amplitude coupling in neuronal network oscillations. Prog. Biophys. Mol. Biol. 105, 49–57. doi: 10.1016/j.pbiomolbio.2010.09.007.

Puig, M. V., and Gener, T. (2015). Serotonin modulation of prefronto-hippocampal rhythms in health and disease. ACS Chem. Neurosci. 6, 1017–1025. doi: 10.1021/cn500350e.

Puig, M. V., and Gulledge, A. T. (2011). Serotonin and prefrontal cortex function: Neurons, networks, and circuits. Mol. Neurobiol. 44, 449–464. doi: 10.1007/s12035-011-8214-0.

Puig, M. V., and Miller, E. K. (2015). Neural substrates of dopamine D2 receptor modulated executive functions in the monkey prefrontal cortex. Cereb. Cortex N. Y. N 1991 25, 2980–2987. doi: 10.1093/cercor/bhu096.

Puig, M. V., Watakabe, A., Ushimaru, M., Yamamori, T., and Kawaguchi, Y. (2010). Serotonin modulates fast-spiking interneuron and synchronous activity in the rat prefrontal cortex through 5-HT1A and 5-HT2A receptors. J. Neurosci. 30, 2211–2222. doi: 10.1523/JNEUROSCI.3335-09.2010.

Rajagopal, L., Massey, B. W., Huang, M., Oyamada, Y., and Meltzer, H. Y. (2014). The novel object *recognition test in rodents in relation to cognitive impairment in schizophrenia*. Curr. Pharm. Des. 20, 5104–5114.

Rajagopal, L., Massey, B. W., Michael, E., and Meltzer, H. Y. (2016). Serotonin (5-HT)1A receptor agonism and 5-HT7 receptor antagonism ameliorate the subchronic phencyclidine-induced deficit in executive functioning in mice. Psychopharmacology (Berl.) 233, 649–660. doi: 10.1007/s00213-015-4137-1.

Sabbagh, J. J., Murtishaw, A. S., Bolton, M. M., Heaney, C. F., Langhardt, M., and Kinney, J. W. (2013). Chronic ketamine produces altered distribution of parvalbumin-positive cells in the hippocampus of adult rats. Neurosci. Lett. 550, 69–74. doi: 10.1016/j.neulet.2013.06.040.

Sigurdsson, T. (2016). Neural circuit dysfunction in schizophrenia: Insights from animal models. Neuroscience 321, 42–65. doi: 10.1016/j.neuroscience.2015.06.059.

Sigurdsson, T., and Duvarci, S. (2016). Hippocampal-prefrontal interactions in cognition, behavior and psychiatric disease. Front. Syst. Neurosci. 9, 190–190. doi: 10.3389/fnsys.2015.00190.

Spanswick, S. C., and Dyck, R. H. (2012). Object/context specific memory deficits following medial frontal cortex damage in mice. PloS One 7, e43698. doi: 10.1371/journal.pone.0043698.

Takehara-Nishiuchi, K. (2020). Prefrontal-hippocampal interaction during the encoding of new memories. Brain Neurosci. Adv. 4, 2398212820925580. doi: 10.1177/2398212820925580.

Tort, A. B. L., Komorowski, R. W., Manns, J. R., Kopell, N. J., and Eichenbaum, H. (2009). Theta-gamma coupling increases during the learning of item-context associations. Proc. Natl. Acad. Sci. 106, 20942–7. doi: 10.1073/pnas.0911331106.

Turetsky, B. I., Dress, E. M., Braff, D. L., Calkins, M. E., Green, M. F., Greenwood, T. A., et al. (2015). The utility of P300 as a schizophrenia endophenotype and predictive biomarker: clinical and socio-demographic modulators in COGS-2. Schizophr. Res. 163, 53–62. doi: 10.1016/j.schres.2014.09.024.

Uhlhaas, P. J., and Singer, W. (2010). Abnormal neural oscillations and synchrony in schizophrenia. Nat. Rev. Neurosci. 11, 100–113. doi: 10.1038/nrn2774.

Umbricht, D., Javitt, D., Novak, G., Bates, J., Pollack, S., Lieberman, J., et al. (1999). Effects of *risperidone on auditory event-related potentials in schizophrenia*. Int. J. Neuropsychopharmacol. 2, 299–304. doi: 10.1017/S1461145799001595.

Vallat, R. (2018). Pingouin: statistics in Python. J. Open Source Softw. 3, 1026. doi: 10.21105/joss.01026.

Vinck, M., Oostenveld, R., Wingerden, M. V., Battaglia, F., and Pennartz, C. M. A. (2011). An improved index of phase-synchronization for electrophysiological data in the presence of volume-conduction, noise and sample-size bias. NeuroImage 55, 1548–1565. doi: 10.1016/j.neuroimage.2011.01.055.

Wang, C., Furlong, T. M., Stratton, P. G., Lee, C. C. Y., Xu, L., Merlin, S., et al. (2021). Hippocampus–prefrontal coupling regulates recognition memory for novelty discrimination. J. Neurosci. 41, 9617–9632. doi: 10.1523/JNEUROSCI.1202-21.2021.

Wang, X., Li, Y., Chen, J., Li, Z., Li, J., and Qin, L. (2020). Aberrant auditory steady-state response of awake mice after single application of the NMDA receptor antagonist MK-801 into the medial geniculate body. Int. J. Neuropsychopharmacol. 23, 459–468. doi: 10.1093/ijnp/pyaa022.

Warburton, E. C., and Brown, M. W. (2015). Neural circuitry for rat recognition memory. Behav. Brain Res. 285, 131–139. doi: 10.1016/j.bbr.2014.09.050.

Zorrilla de San Martín, J., Donato, C., Peixoto, J., Aguirre, A., Choudhary, V., De Stasi, A. M., et al. (2020). Alterations of specific cortical GABAergic circuits underlie abnormal network activity in a mouse model of Down syndrome. eLife 9, e58731. doi: 10.7554/eLife.58731.

