## Supplementary Information for "Neural substrates of cognitive impairment in a NMDAR hypofunction mouse model of schizophrenia and rescue by risperidone"

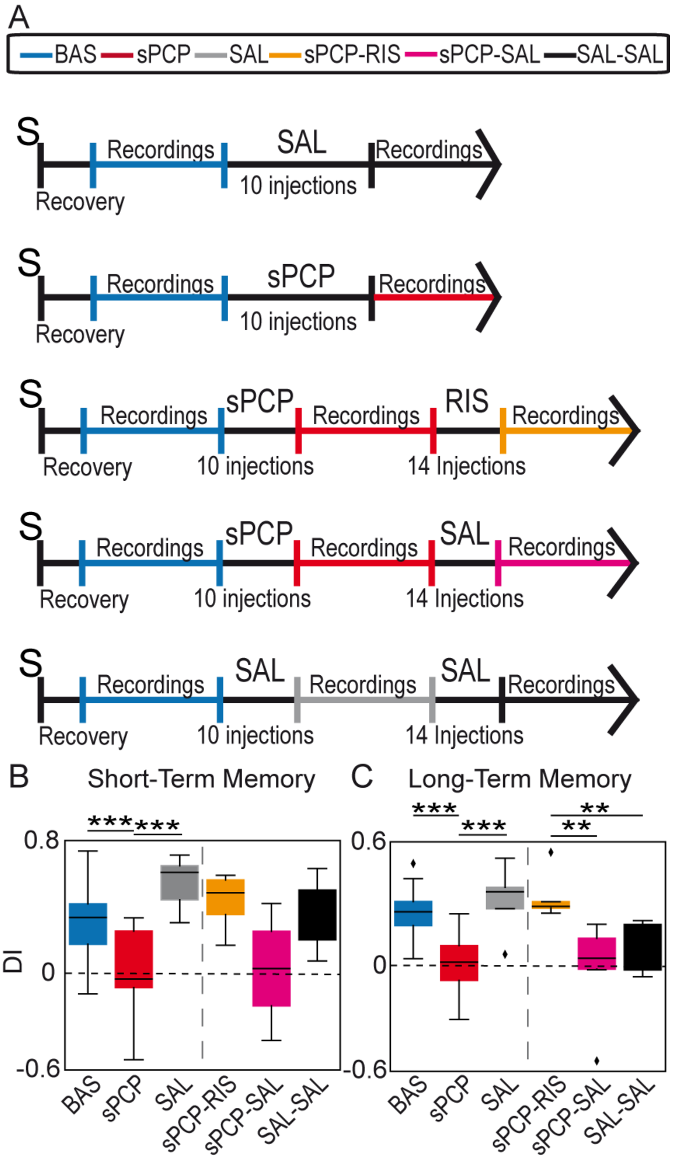


**Supplementary Figure 1: (A)** Experimental groups of the study: sPCP (*n* = 21 mice), SAL (*n* = 7), sPCP-RIS (*n* = 9), sPCP-SAL (*n* = 7) and SAL-SAL (*n* = 7). **(B)** Discrimination indices of all the pharmacological groups investigated during the 3-minute (short-term) memory test. BAS vs. sPCP vs. SAL: $F_{(1,20)} = 26.31$, $p < 0.0005$, one-way ANOVA. sPCP-RIS vs. sPCP-SAL vs. SAL-SAL: $F_{(2,24)} = 5.11$, $p = 0.017$, two-way ANOVA. **(C)** Discrimination indices for all the pharmacological groups investigated during the 24h (long-term) memory test. BAS vs. sPCP vs. SAL: $F_{(1,26)} = 15.26$, $p = 0.0006$, one-way ANOVA. sPCP-RIS vs. sPCP-SAL vs. SAL-SAL: $F_{(2,24)} = 15.76$, $p = < 0.0005$, two-way ANOVA. The S indicates the day of surgery. Vertical dashed lines separate the two statistical groups.

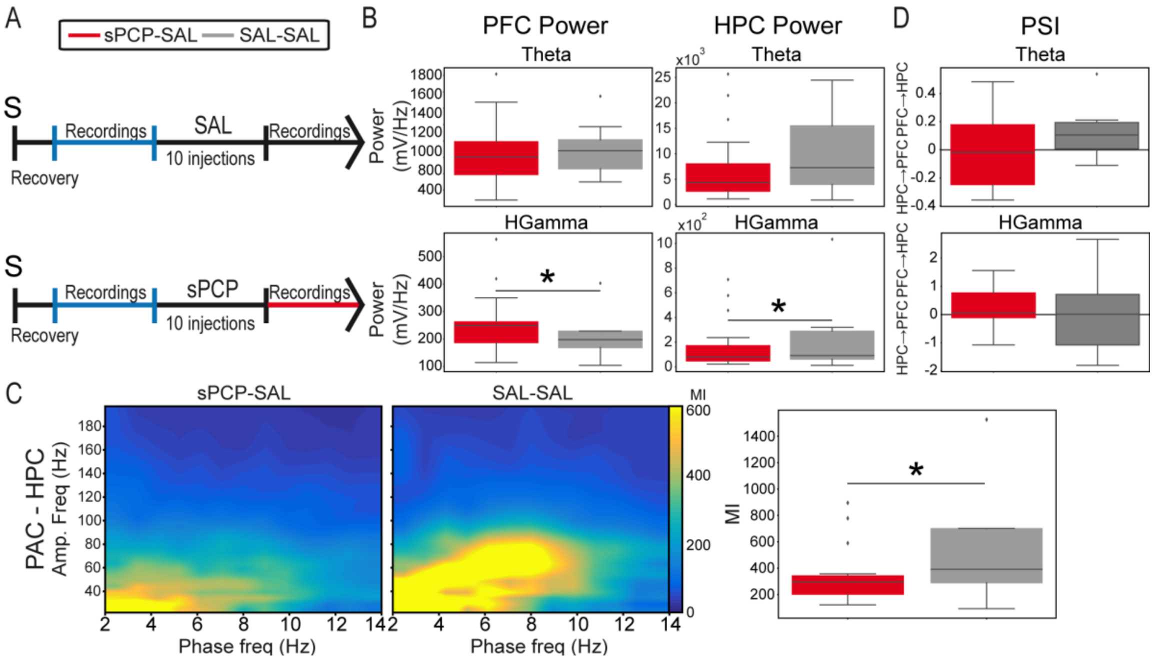


**Supplementary Figure 2:** Saline controls (SAL group) did not exhibit the alterations observed in the sPCP group during quiet wakefulness. **(A)** Experimental protocols of the sPCP and SAL groups. **(B)** Quantification of theta and high gamma power in both groups. High gamma power increased in the PFC and decreased in the HPC after sPCP but not after saline (sPCP vs. SAL: $F_{(1,22)} = 6.59$, $p = 0.016$; two-way ANOVA). **(C)** Local and inter-regional theta-gamma coupling weakened after sPCP but not after saline (sPCP vs. SAL: $F_{(1,22)} = 6.31, 4.62$, $p = 0.02, 0.024$; two-way ANOVA). **(D)** The directionality of signals within prefrontal-hippocampal circuits was not affected by sPCP or saline. The S indicates the day of surgery.

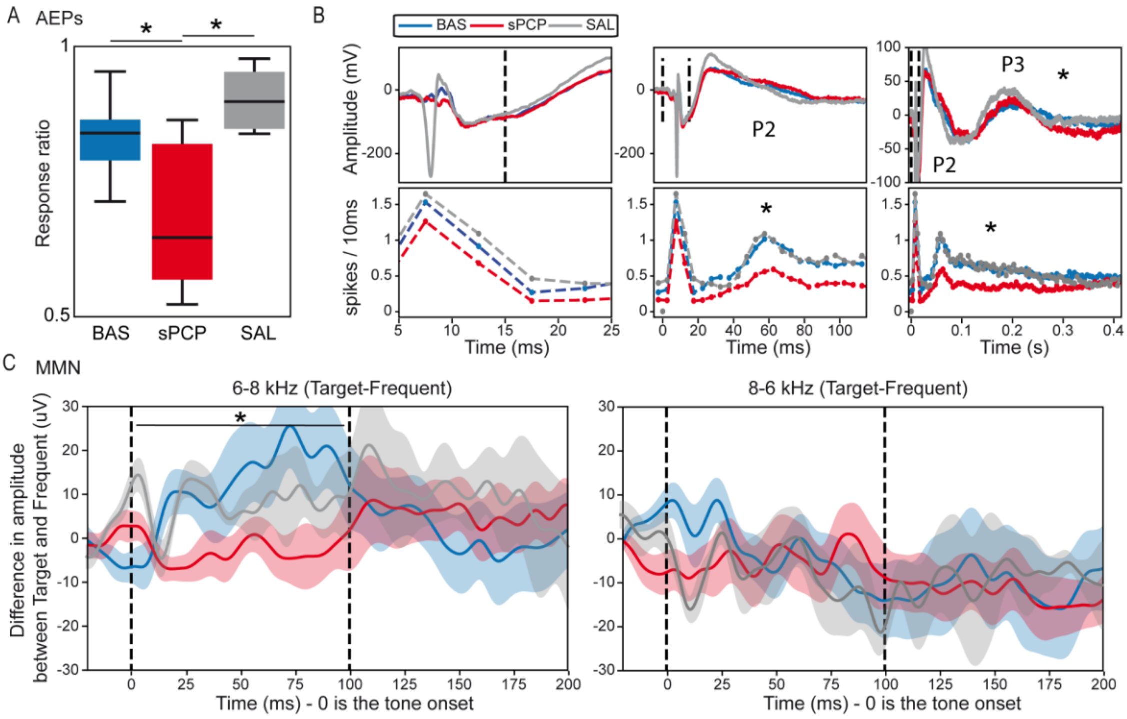


**Supplementary Figure 3:** After the administration of saline (SAL control group) the behavioural and neurophysiological fingerprints of auditory processing were similar to baseline. **(A)** Ratio of auditory evoked potential (AEPs) responses detected in the PFC after the presentation of 100 auditory stimuli. The response ratio remained elevated after saline, but not after sPCP. **(B)** AEPs and corresponding spiking activity (multi-unit firing rates) in the PFC at three different timescales. The neuro-physiological responses were very similar between baseline and after saline. **(C)** Mismatch negativity (MMN) was detected during the presentation of the 6-8KHz target-frequent tone combination. MMN was present in the saline-treated but not the sPCP-treated group. Vertical dashed lines mark the start and end of tone presentation.
